# Spontaneous Alignment of Myotubes through Myogenic Progenitor Cell Migration

**DOI:** 10.1101/2023.07.10.548397

**Authors:** Lauren E. Mehanna, Adrianna R. Osborne, Charlotte A. Peterson, Brad J. Berron

## Abstract

In large volume muscle injuries, widespread damage to muscle fibers and the surrounding connective tissue prevents myogenic progenitor cells (MPCs) from initiating repair. There is a clinical need to rapidly fabricate large muscle tissue constructs for integration at the site of large volume muscle injuries. Most strategies for myotube alignment require microfabricated structures or prolonged orientation times. We utilize the MPC’s natural propensity to close gaps across an injury site to guide alignment on collagen I. When MPCs are exposed to an open boundary free of cells, they migrate unidirectionally into the cell-free region and align perpendicular to the original boundary direction. We study the utility of this phenomenon with biotin - streptavidin adhesion to position the cells on the substrate, and then demonstrate the robustness of this strategy with unmodified cells, creating a promising tool for MPC patterning without interrupting their natural function. We pre-position MPCs in straight-line patterns separated with small gaps. This temporary positioning initiates the migratory nature of the MPCs to align and form myotubes across the gaps, similar to how they migrate and align with a single open boundary. There is a directional component to the MPC migration perpendicular (90°) to the original biotin-streptavidin surface patterns. The expression of myosin heavy chain, the motor protein of muscle thick filaments, is confirmed through immunocytochemistry (ICC) in myotubes generated from MPCs in our patterning process, acting as a marker of skeletal muscle differentiation. The rapid and highly specific binding of biotin-streptavidin allows for quick formation of temporary patterns, with MPC alignment based on natural regenerative behavior rather than complex fabrication techniques.

**Impact Statement:** Positioning myogenic progenitor cells (MPCs) into straight-line patterns with intentional spacings initiates the migration of these cells to bridge these gaps, mimicking their behavior in response to small-scale injuries. By creating repetitions of patterned cells and spacings, we have demonstrated rapid migration and alignment of MPCs, which differentiate into a long-range 2D layer of aligned myotubes.

## 1. Introduction

Despite the intrinsic regenerative capacity of muscle tissue, large skeletal muscle injuries typically lead to functional irregularities (1, 2). Large skeletal muscle injuries from surgical losses, extreme exercise, car accidents, or military traumas (3–5) are challenging because they create a pro-inflammatory microenvironment that overwhelms the body’s innate repair mechanism (6, 7). Damage to greater than 20% damage to a given muscle’s mass is categorized as volumetric muscle loss (VML) (3, 8), and the current standard of care for these injuries is physical therapy or autologous tissue transfer. Even with prolonged physical therapy, persistent strength deficits remain, and only partial range of motion is restored to the remaining tissue (9–11). Surgical complications limit the success achieved through autologous tissue transfer, as infection can result in total graft rejection or donor site morbidity can occur due to tissue necrosis (12–14). When tissue transfer does occur, typically only limited blood flow and active motion are restored (14).

In the absence of autologous tissue options, new generations of engineered tissue constructs seek to recreate highly organized and aligned muscle fibers in the native tissue architecture prior to reinsertion at the injury (15). These constructs must be integrated prior to the fibrotic response beginning at 1-2 weeks post injury, otherwise fibrosis will limit complete restoration of tissue function (16).

Engineered scaffolds for VML repair utilize quiescent muscle stem cells, called satellite cells, to form functional, in vitro myotube structures. In small volume muscle injuries, these satellite cells activate as myogenic progenitor cells (MPCs), travel through basal lamina to the injury site, and proliferate in response to inflammatory signals to restore the injured tissue (17). MPCs work to bridge the gap over the injury and with time fuse together to form multinucleated myotube structures (3, 18). The innate tube formation capacity of MPCs in vitro is the foundation of most modern VML tissue engineering strategies.

Many tissue construct designs for VML start as two-dimensional layers of aligned myotube structures prior to being scaled in three dimensions. Current designs predominantly seed MPCs on a substrate material comprised of one or more extracellular matrix (ECM) proteins (19).

Collagen I is the main structural component in skeletal muscle ECM and is an ideal substrate for MPC seeding and alignment. Current strategies for myotube alignment primarily alter and align the substrate itself to guide MPC placement and orientation into patterned structures. Research studies using lithography (15, 20, 21), electrospinning (22–24), 3D bioprinting (25–27), shear-based extrusion (28), and rolling (29) are currently employed to engineer the scaffold and create highly aligned structures. While these approaches have been successful in achieving myotube formation with aligned orientation, they can be expensive, complex to fabricate, and have a lower size limit of the patterns that can be achieved (30). Instead, we take advantage of the natural migratory behavior of MPCs to create rapid alignment of myotube structures.

In this work, MPC alignment is based on natural migration and regenerative behavior rather than complex mechanical or topographical cues. We utilize the MPC’s natural propensity to close gaps across an injury site to guide alignment on a collagen I protein substrate. Through the binding affinity of biotin and streptavidin molecules, known for forming strong non-covalent interactions (31, 32), we pre-position MPCs in straight-line patterns separated with small gaps, mimicking many small-scale muscle injuries. Functionalized biotin groups bind to primary amine groups present on both the collagen I surface and amino acids in the cell membrane (33). Because streptavidin has four binding sites, it is capable of binding to multiple biotin molecules, acting as a linker to aid specific placement of biotinylated cells wherever it is bound to the patterned biotin on the substrate surface. This biotin-streptavidin cell patterning method is desirable as it is rapid, inexpensive, uses commercially available reagents, and allows for pattern customization based on the photomask design. The temporary positioning of MPCs using biotin-streptavidin conjugations initiates their migratory nature to bridge the gap as they would after an injury, aligning in one direction over the gap and forming myotubes structures (Fig. 1) (3).

**Figure 1.**
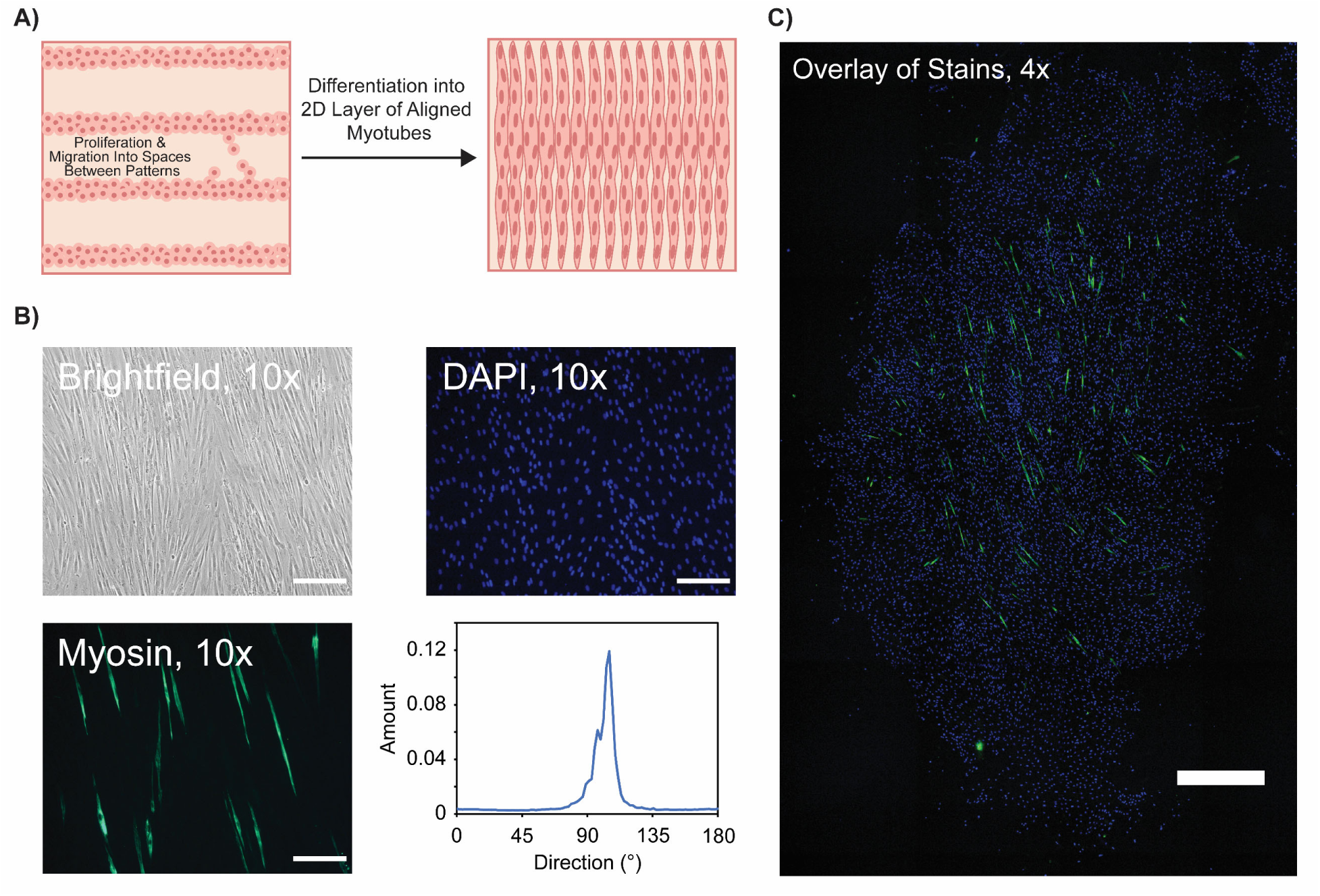
Large-scale alignment of myotubes after positioning MPCs in patterned structures. A) Schematic representation of MPCs prep-positioned on the substrate in straight-line patterns. During differentiation, these cells migrate into the spacings between patterns and form myotubes perpendicular to the original patterns. B) Representative images (brightfield, MyHC, DAPI) of aligned myotubes formed after patterned MPCs were placed in differentiation media for 6 days. Myotube alignment was characterized using the ImageJ Directionality plug-in for MyHC fluorescent images. Imaged were taken using brightfield and fluorescent microscopy at 10x magnification. C) Full-well image (1.7 cm^2^ patterned area) taken at 4x magnification overlaying MyHC and DAPI staining of aligned myotubes formed after patterned MPCs were placed in differentiation media for 6 days. Scale bars represent 200 µm in B and 1000 μm in C.

## 2. Materials and Methods

### 2.1 Cell Culture

Human primary myogenic progenitor cells (MPCs) were obtained from the healthy muscle biorepository in the Center for Muscle Biology, University of Kentucky and from the National Disease Research Exchange (NDRI, Res Code DPEC6 001). MPCs were cultured at 37°C and 5% CO_2_ in Hams F-12 Nutrient Mixture (Thermo Fisher, Waltham, MA) supplemented with 20% FBS (Biotechne, Minneapolis, MN), 1% penicillin-streptomycin (pen-strep) (HyClone, Logan, UT), and 1% antibiotic-antimycotic. A stock solution of 100 µg basic fibroblast growth factor (bFGF) (Peprotech, Rocky Hill, NJ) was dissolved with 5 mg bovine serum albumin (BSA) (Millipore Sigma, Burlington, MA) and 1 mL phosphate buffered saline (PBS) (Millipore Sigma, Burlington, MA). The growth media was supplemented with 1 µL of the bFGF stock solution per 1 mL media. MPCs were cultured in 75 cm^2^ tissue culture flasks (VWR, Radnor, PA) to 20-50% confluency and passaged using 10x Trypsin-EDTA (Thermo Fisher, Waltham, MA) diluted to 1x concentration in PBS. During patterning experiments, MPCs were seeded on Biocoat collagen I coated 4-well slides (Corning, Corning, NY). All solutions throughout the patterning process were prepared in 1x PBS supplemented with 1% glucose and 1% penicillin-streptomycin.

### 2.2 Directional Migration Analysis Through Cell Exclusion

A 2-well silicone insert (Ibidi, Fitchburg, WI) with a defined 500 µm cell free gap between the wells was placed on a collagen I slide. Two controls (negative control and adhesive control) of MPCs were seeded at a density of 2.27 × 10^5^ cells/cm^2^ in differentiation media consisting of DMEM supplemented with 2% Horse Serum (Atlanta Biologicals, Flowery Branch, GA), 1% pen-strep, and 1% antibiotic-antimycotic in the 2 wells of the insert and incubated at 37° C and 5% CO_2_ for 2 h to allow for cell adherence. The negative control of unmodified MPCs was seeded on unmodified collagen I and the adhesive control of biotinylated MPCs was seeded on a collagen I surface modified non-specifically (no patterns) with TFPA-PEG3-Biotin and Streptavidin-Cy3. After 2 h, the culture inserts were removed, and the surface was washed with fresh media to remove any unadhered cells. The slides were replenished with fresh media and the wells were imaged using brightfield microscopy to observe the 500 µm cell free gap between the regions of adhered cells as well as the edges of the cell area. The slides were incubated at 37° C and 5% CO_2_ and imaged every 24 h for 48 h to observe cell migration into the cell-free regions.

The direction and alignment of cell migration was analyzed in the original cell-free regions only using the ImageJ (NIH) Directionality plug-in, with the Fourier components method using 90 bins between 0 and 180° to track vertical cell migration and alignment. A direction of 0° represented a horizontal orientation while a direction of 90° represented a vertical orientation. ImageJ output a histogram with relative alignment over a range of directions and applied a Gaussian fit to the histogram. The ‘Direction’ reported by ImageJ represented the center of the Gaussian while the ‘Dispersion’ represented the standard deviation of the Gaussian (Direction ± Dispersion). The ‘Amount’ represented the number of structures in a given direction, which was calculated as the sum of the center – standard deviation to center + standard deviation, divided by the total sum of the histogram.

### 2.3 Substrate Patterning with Biotin-Streptavidin

The substrate of interest was coated with 5 mg/mL BSA for 1 h to add a non-specific protein layer with free amine groups to the slide surface, then washed with Deionized (DI) water and air dried. A TFPA-PEG3-Biotin (Thermo Fisher, Waltham, MA) stock solution was prepared by dissolving 25 mg in DMSO (VWR, Radnor, PA) to a concentration of 100 mg/mL. The stock solution was diluted with PBS to create a 1 mg/mL working solution. 20 μL of the working solution was added to the patterned region on the given slide, then covered with a chrome photomask etched with the desired pattern size (University of Louisville Micro/Nano Technology Center, KY). The sample covered with the photomask was exposed to a 365 nm UV light (Thor Labs, Newton, NJ) with 1 mW/cm^2^ intensity for 5 min. The sample and photomask were each washed with DI water and air dried. For adhesive controls without patterned structures, the working TFPA-PEG3-Biotin solution was added to the slide and irradiated without the photomask to allow binding to the entire surface. After the surface was photopatterned with biotin, 20 μL/mL Streptavidin-Cy3 (Invitrogen, Waltham, MA) was added to the well for 30 min. The sample was rinsed again with DI water and air dried. The biotin-streptavidin conjugations on the substrate surface were verified using fluorescent microscopy on an Eclipse Ti-U inverted microscope (Nikon, Minato City, Tokyo, Japan). Pattern sizes of 50 μm,100 μm, and 200 μm line widths with 800 μm spacing between lines were studied. To compare to the negative and adhesive control samples seeded in the 2-well silicone inserts, additional samples were also patterned with only half of the wells irradiated to contain the biotin-streptavidin conjugations while the other half of the well remained unmodified.

### 2.4 Cell Surface Modification with Biotin

After being trypsinized, MPCs were washed twice for 3 min at 400 RCF at room temperature with PBS supplemented with 1% glucose and 1% pen-strep (Eppendorf 5702R, Hamburg, Germany). Biotinylation of the cell surface was performed by adding 1 mM Sulfo-NHS-LC-Biotin (Thermo Fisher, Waltham, MA) to the cell pellet at 250 μL per 1 million cells. The cells were incubated for 40 min, then washed twice again at 3 min at 400 RCF. The cell pellet was resuspended in PBS with 1% glucose and 1% pen-strep and the cells were seeded on the patterned well on the slide for 40 min in darkness by covering with aluminum foil. MPCs were seeded on a 1.7 cm^2^ collagen I well at a seeding density of 1.18 × 10^5^ cells/cm^2^. Cell patterned slides were washed using a micropipette with PBS to maximize cell adhesion in the patterned regions while minimizing cell adhesion in the spaces between patterns. Once washed, the slides were placed in a four-well rectangular dish (Nunc, Thermo Fisher, Waltham, MA) fully submerged cell-side up with 5 mL fresh PBS with 1% glucose and 1% pen-strep so that cell patterns could be easily imaged.

**Figure 2.**
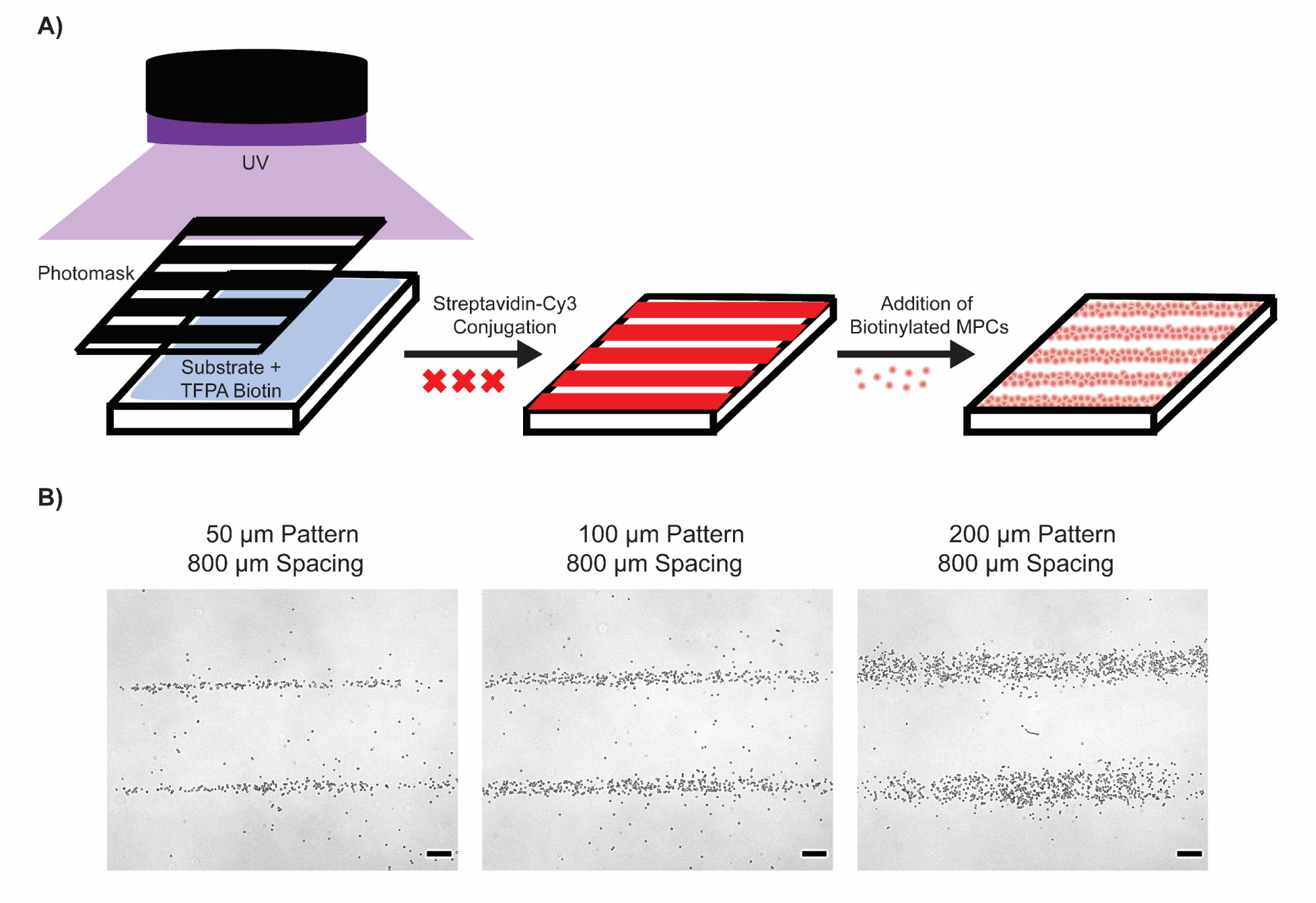
MPC patterning using biotin-streptavidin conjugations. A) Schematic representation of the patterning process of MPCs on collagen I using biotin-streptavidin conjugations. B) Representative images of MPCs patterned in 50, 100, and 200 μm line width patterns with 800 μm spacings between lines. Patterns were imaged using brightfield microscopy and 4x magnification. Scale bars represent 200 μm.

### 2.5 Differentiation and Staining of Myotubes

Once patterned and imaged, samples were placed into differentiation media and incubated for 6 days at 37° C and 5% CO_2_, with the media changed every 2 days. Samples were imaged every two days to monitor myotube formation and cell alignment. Cell alignment was analyzed using ImageJ (NIH) Directionality plug-in, as described in Section 2.2. Additionally, the same control conditions of MPCs (negative control and adhesive control) as described in Section 2.2 were seeded at 1.18 × 10^5^ cells/cm^2^ on the entirety of the collagen slide (no insert) in differentiation media. The negative control of unmodified MPCs was seeded on unmodified collagen I and the adhesive control of biotinylated MPCs was seeded on a collagen I surface modified non-specifically (no patterns) with TFPA-PEG3-Biotin and Streptavidin-Cy3. On day 6, all slides (patterns and controls) were fixed with 4% formaldehyde (Millipore Sigma, Burlington, MA) for 20 min. The slides were washed three times with PBS, then incubated in a permeabilization solution for 30 min. The permeabilization solution contained 20 mM citrate phosphate (Millipore Sigma, Burlington, MA), 0.1 mM EDTA (Millipore Sigma, Burlington, MA), 0.2 mM sucrose (Millipore Sigma, Burlington, MA), 0.1% Triton X-100 (Millipore Sigma, Burlington, MA), diluted in DI water to make a 50 mL total solution. The slides were washed three times with PBS again, then incubated overnight at 4° C with a pan myosin heavy chain (MyHC) primary antibody (DSHB A4.1025, Iowa City, IA) in a moist chamber on a rocker. Samples were washed once with the permeabilization solution, followed by two washes with PBS. A 1:200 goat, anti-mouse IgG2a Alexa Fluor 488 (Invitrogen, Waltham, MA) secondary antibody in TBS-T was added to the samples and incubated for 1 h on a rocker. The samples were washed three times with PBS. A 1:10,000 DAPI nuclear stain in PBS was added to the samples for an additional 20 min on a rocker, followed by three washes with PBS. The samples were then placed in fresh PBS and imaged using the Eclipse Ti-U inverted microscope (Nikon, Minato City, Tokyo, Japan) to observe myotube formation. The entire 1.7 cm^2^ patterned area of the slide was also imaged using the large scan tiling feature on the Eclipse Ti-E (Nikon, Minato City, Tokyo, Japan) inverted microscope.

## 3. Results

### 3.1 Directional Cell Migration through Cell Exclusion

Cell migration was studied through a cell exclusion assay, in which cells were seeded in two individual wells with a barrier between the two. Once the cells were adhered to the bottom of the slide, the wells were removed, revealing a cell-free gap between the two wells of cells and a large cell-free area on the top and bottom boundaries of each well. Cell migration was first studied at the top boundary of each well, in which only one directional migration is possible. The orientation of cells was only analyzed in the cell-free region above where the silicone insert was placed (above blue line) in the first two days after seeding using the ImageJ Directionality plugin. Representative images of each condition (negative control, adhesive control, and patterned alignment) are shown in Fig. 3.

**Figure 3.**
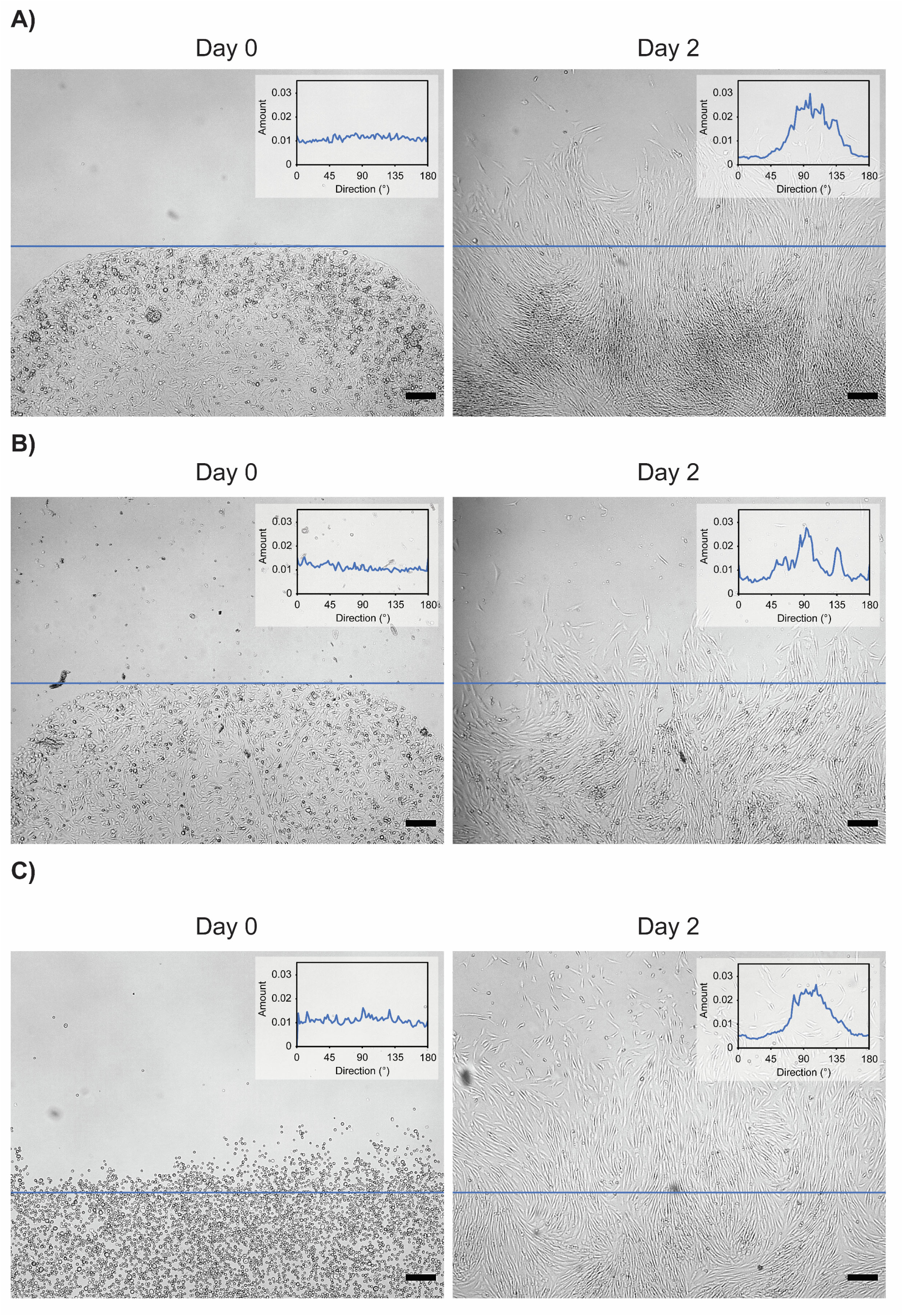
Cell migration analysis of MPCs with open boundary. A) Representative images and directionality analysis of unmodified MPCs seeded on an unmodified collagen I surface (negative control) in Ibidi silicone inserts. Images were taken at the top boundary of the well directly after the insert was removed on day 0 and again on day 2. Directionality analysis was completed above the blue line, which indicates the original cell-free region outside of the insert boundary. B) Representative images and directionality analysis of biotinylated MPCs seeded on a collagen I surface modified non-specifically with TFPA-PEG3-Biotin and Streptavidin-Cy3 (adhesive control) in Ibidi silicone inserts. Images were taken at the top boundary of the well directly after the insert was removed on day 0 and again on day 2. Directionality analysis was completed above the blue line, which indicates the original cell-free region outside of the insert boundary. C) Representative images and directionality analysis of biotinylated MPCs patterned on half of the collagen surface using biotin-streptavidin positioning without Ibidi silicone inserts. Images were taken at the top boundary of the well directly after the insert was removed on day 0 and again on day 2. Directionality analysis was completed above the blue line, which indicates the boundary of biotin-streptavidin conjugations that guided cell seeding. All images were taken using brightfield microscopy at 4x magnification. Scale bars represent 200 µm.

There was a clear visual migratory response to cells on the boundary (blue line) for any condition within the first two days in differentiation media, as shown qualitatively in the brightfield images. There was no clear directional orientation of cells on day 0, but cells aligned perpendicular to the boundary into the cell-free space by day 2 in all seeding conditions, indicated by the clear peak formation in the day 2 directional graphs. MPCs on the negative control had no established directionality on day 0. However, by day 2, a direction of 102° ± 28° was visible, with at least 88% of cells in this orientation. Similar MPC alignment was seen with the adhesive control, with at least 37% of cells having an established directionality of 92° ± 10° on day 2. When MPCs were patterned using the biotin-streptavidin cell patterning approach to target cell placement on half of the slide well, no directionality was clear visually on day 0, but at least 77% of cells were oriented in the 101° ± 24° direction on day 2.

When the exclusion assays were imaged with a single spacing between the two wells of cells (marked with blue), the MPCs still migrated into the cell-free space between the wells. Within the first two days of the wells being removed, the cells had completely migrated into the original cell-free regions and aligned perpendicular to the original horizontal well direction. The directionality of the cells was only analyzed in the spacing between the original wells (region between blue lines) to determine the cell response when exposed to an intentional spacing free of other cells (Fig. 4).

**Figure 4.**
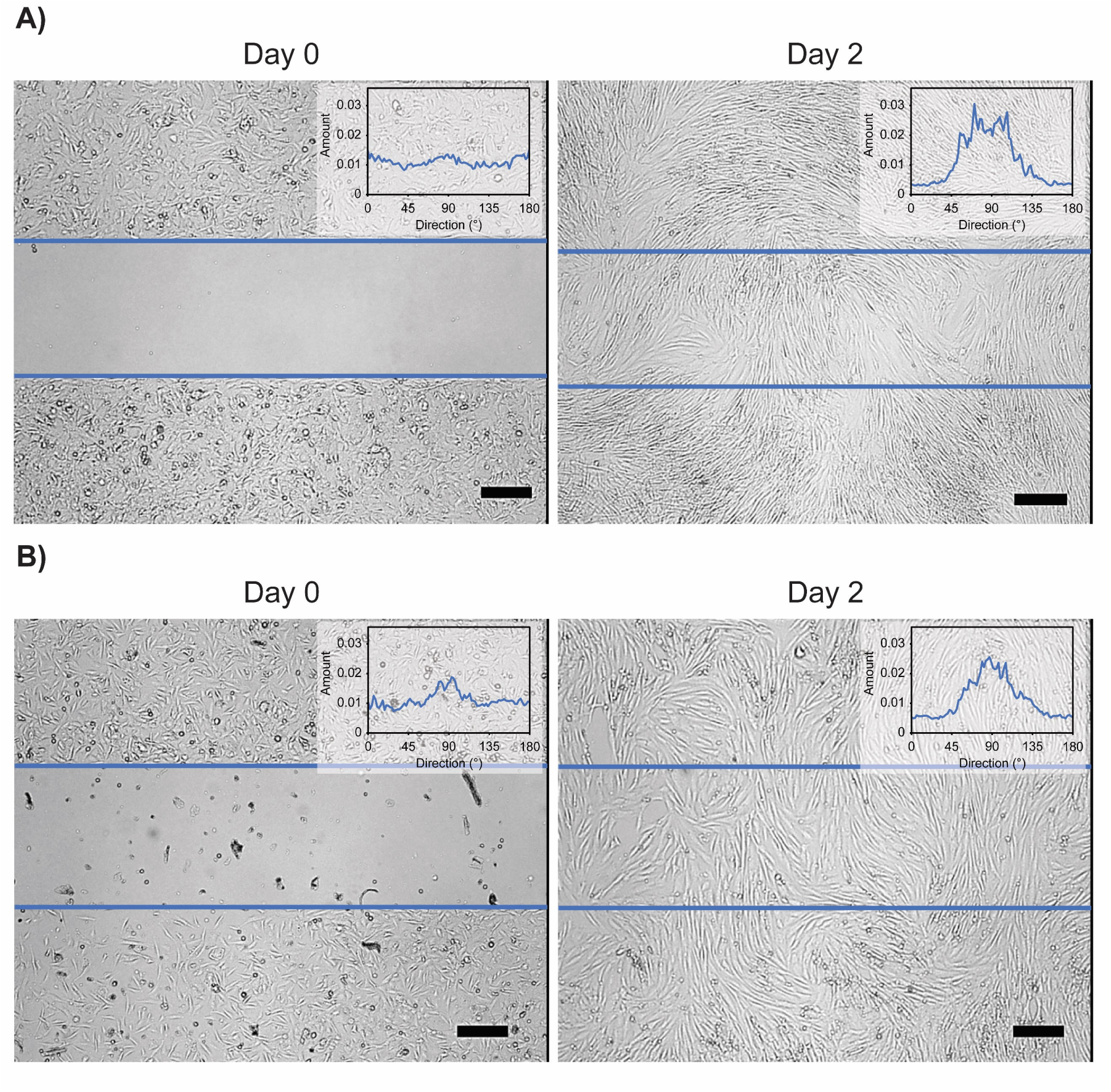
Cell migration analysis of MPCs between wells with intentional spacing. A) Representative images and directionality analysis of unmodified MPCs seeded on an unmodified collagen I surface (negative control) in Ibidi silicone inserts. Images were taken in the spacing between the wells directly after the insert was removed on day 0 and again on day 2. B) Representative images and directionality analysis of biotinylated MPCs seeded on a collagen I surface modified non-specifically with TFPA-PEG3-Biotin and Streptavidin-Cy3 (adhesive control) in Ibidi silicone inserts. Images were taken in the spacing between the wells directly after the insert was removed on day 0 and again on day 2. All images were taken using brightfield microscopy at 4x magnification. Scale bars represent 200 µm.

MPCs seeded on both the negative control and adhesive control had no established directionality in the cell-free region on day 0. By day 2, visible cellular orientation was established for both conditions. The negative control had at least 86% of cells oriented in the direction of 86° ± 27° and the adhesive control had at least 75% of cells oriented in the direction of 90° ± 24°.

### 3.3 Directional Migration of MPCs Patterned in Repeating Structures with Intentional Spacings

Using the biotin-streptavidin patterning approach, cells were placed in repeating temporary straight-line structures of 200µm thickness with intentional spacings of 800 µm. The patterns were placed in differentiation media and imaged every 2 days to observe cell migration and orientation. Within the first 2 days, the cells migrated from the original pattern positions into the cell-free spacings, similar to the cell exclusion studies with a single spacing, with the original cell patterns no longer noticeable through brightfield imaging by day 2 (Fig. 5).

**Figure 5.**
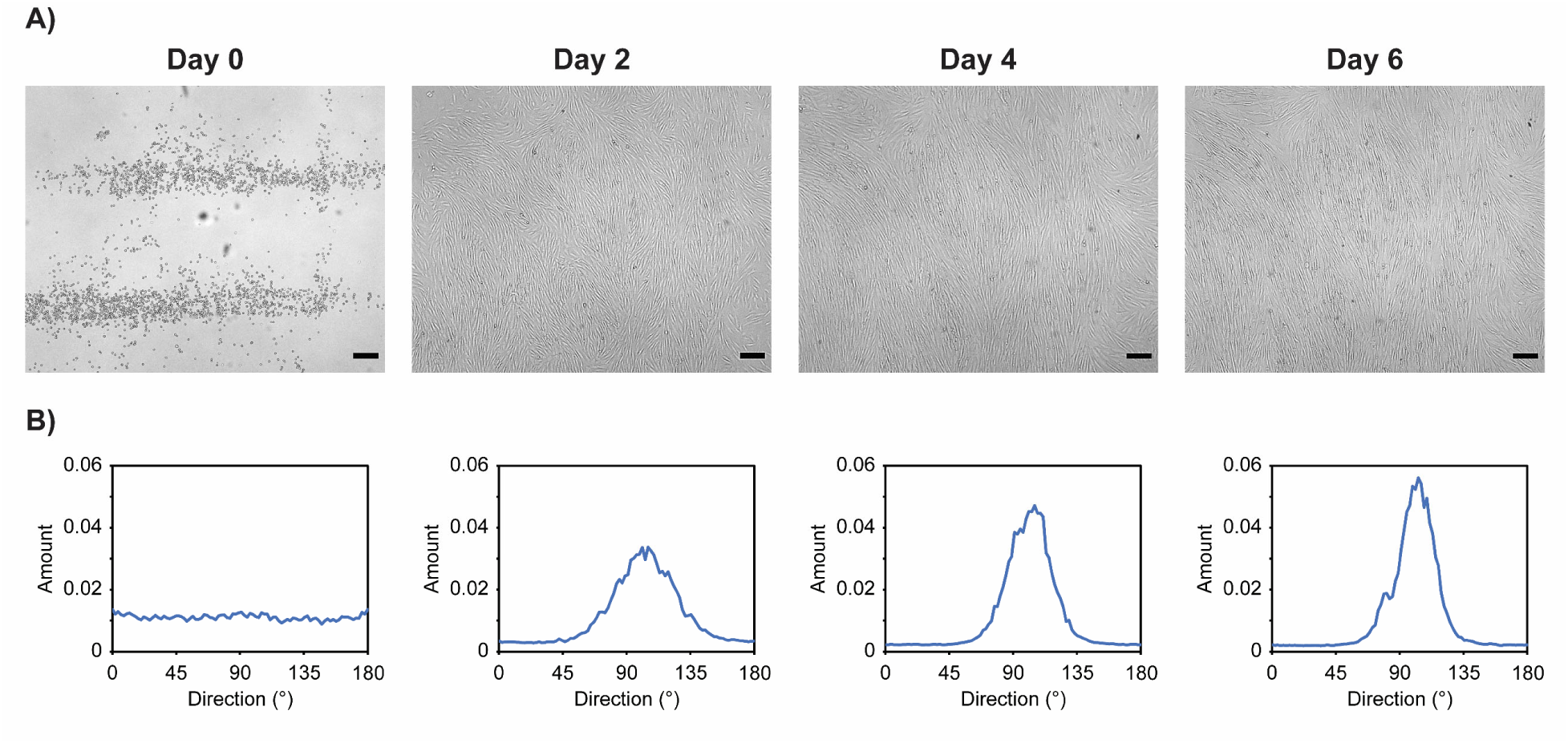
Cell migration analysis of MPCs patterned with biotin-streptavidin. A) Representative images of MPCs placed in temporary patterns of 200 µm line widths with 800 µm spacings using biotin-streptavidin conjugations on day 0 and observed in differentiation media every 2 days for 6 days. All images were taken using brightfield microscopy at 4x magnification. Scale bars represent 200 µm. B) Directionality analysis of MPCs at each timepoint mentioned in A.

Cells migrated and aligned perpendicular to the original patterns. There was no clear cell orientation established on day 0. By day 2 a visible directionality peak was established with at least 80% of cells in the direction of 103° ± 20°. This directionality remained similar over the next 4 days, with 82% of cells in a direction of 102° ± 15° on day 4 and 79% of cells in a direction of 102° ± 12° on day 6 as the MPCs formed myotube structures and continued to mature.

### 3.4 MPC Function with Biotin-Streptavidin Conjugations Analyzed through Tube Formation

MPCs naturally form myotube structures when placed in differentiation media over a 6-day period. We related the functionality of MPCs when modified and unmodified with biotin by studying their ability to form myotubes in each condition previously mentioned (negative control and pattern alignment) (Fig. 6).

**Figure 6.**
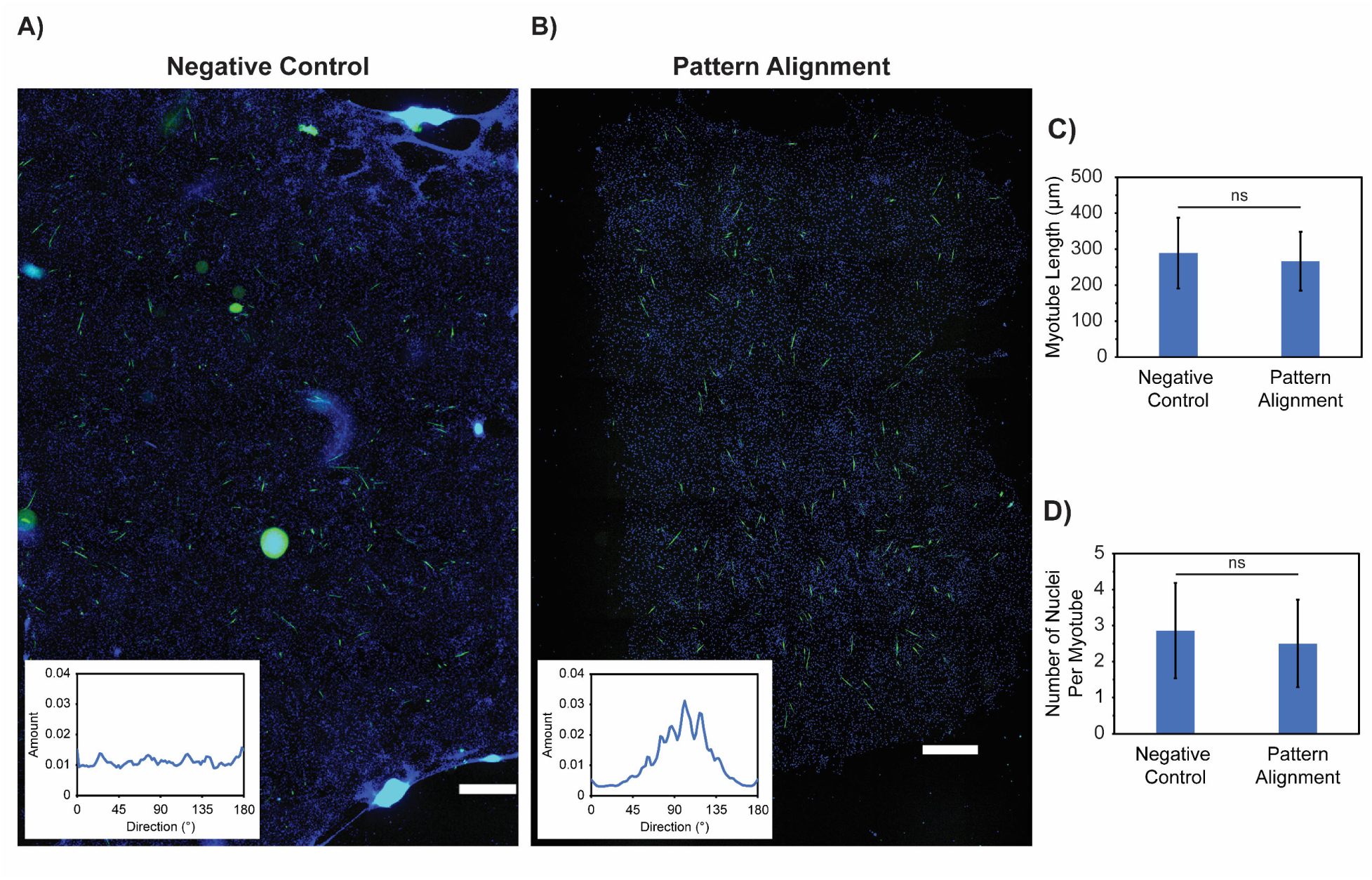
MPC functionality assessed through myotube formation. Representative full-well images and directionality analysis of myotubes formed from MPCs seeded in each condition. A) The negative control represents MPCs seeded on an unmodified collagen I surface. B) The pattern alignment represents biotinylated MPCs seeded in 200 µm patterns with 800 µm spacing. Samples were fixed on day 6 and stained with anti-myosin (green) and DAPI (blue), then imaged using fluorescent microscopy at 4x magnification. Directionality analysis was performed on day 6 full-well myosin staining (green). Scale bars represent 1000 µm. C) Quantitative analysis of the number of nuclei per myotube for each condition described in A and B. Data is shown as mean ± standard deviation of n = 50 myotubes across 3 independent samples for each condition. Significance testing compared each condition using a two-group independent t-test (* p < 0.05, “ns” means not significant). D) Quantitative analysis of myotube length for each condition described in A and B. Data is shown as mean ± standard deviation of n = 50 myotubes across 3 independent samples for each condition. Significance testing compared each condition using a two-group independent t-test (* p < 0.05, “ns” means not significant).

There was a clear directional response when comparing the day 6 myosin staining in the full-well fluorescent images (Fig. 6A, green) for myotubes formed from patterned MPCs versus myotubes formed from the negative control condition. There was no established orientation for myotubes formed in the negative control condition, with only 7% of cells oriented in any given direction. However, there was a visible directionality peak for myotubes patterned using biotin-streptavidin conjugations, with at least 84% of cells oriented in the direction of 101° ± 26°.

There was no significant difference in the length of myotubes formed within any of the seeding conditions (Supplementary Table 1). The negative control had an average myotube length of 289 µm, while patterned MPCs had an average myotube length 267 µm (p = 0.47). Additionally, there was not a significant difference in the number of nuclei per myotube between the negative control with the patterned MPCs (Supplementary Table 1). The negative control had on average 2.86 nuclei per myotube, while the patterned MPCs had on average 2.50 nuclei per myotube (p = 0.53).

## 4. Discussion

Skeletal muscle regeneration is driven by the activation of MPCs following injury (14). When a small-scale injury occurs, MPCs respond by migrating and proliferating to bridge the gap across the injury site, then differentiating to create new myotube structures (3). The goal of this study was to understand the potential role of MPC migration in the alignment of MPCs and the newly formed myotubes in vitro.

We first studied the MPC response to a large cell-free region in one direction by using a cell exclusion assay. When MPCs were exposed to a large area free of cells, the cells unidirectionally migrated into the open space and oriented perpendicular to the original boundary direction (Fig. 3). This supports the hypothesis that MPCs migratory behavior is activated when free regions of substrate are present, similar to their nature in culture as they spread and connect with nearby cells to form tube structures rather than aggregate. This cell migration and alignment was observed by both unmodified MPCs seeded on an unmodified collagen I surface and by biotinylated MPCs seeded on a collagen I surface conjugated with biotin-streptavidin (negative control, adhesive control, pattern alignment). Using biotin-streptavidin conjugations to adhere the cells to the surface did not impact their ability to migrate or align, achieving similar directionality to MPCs seeded directly on unmodified collagen. Instead, these conjugations only created a temporary positioning of the cells on the substrate, which did not alter their natural migratory behavior or overall function. The biotin-streptavidin patterned structures diminished over time in cell culture conditions, based on tracking with fluorescent intensity (Supplementary Fig. 2). Therefore, biotin-streptavidin conjugations are now used to create more advanced architectures of MPCs in vitro. When MPCs were placed in a single circular feature using biotin-streptavidin positioning with exposed substrate on all sides, the cells outwardly migrated radially (Supplementary Fig. 1). Cell directionality was observed perpendicular to the original border around the circular feature. This reinforces that MPCs will migrate and align in any direction of cell-free substrate.

Knowing that MPCs migrate in response to the absence of cells on the substrate, we studied how cells would migrate when exposed to a linear gap rather than having a large open boundary. The MPCs now migrated into the spacing from both regions where the cells were seeded until the spacing was fully closed, mimicking the exact MPC behavior when they respond to injury and regenerate new myotube structures. This single-gap system indicates that small-range ordering is possible and that MPCs will orient in the spacing perpendicular to the original seeding boundary (Fig. 4). Biotin-streptavidin conjugations again did not inhibit the MPC migration or alignment, therefore not hindering their natural function.

In studies with only one spacing present between two large regions of cells, cell orientation was only established in the region of the spacing itself; however, long-range order was not imposed on the larger regions where the cells were originally seeded. With the goal of creating large scale aligned muscle tissues in vitro, we created repetitive gaps to extend small scale MPC alignment strategies to macro scale myotube alignment. By using UV-initiated TFPA biotin with a photomask etched with customized features, we tailored the specific straight-line patterns for the cells to bind. Creating repeating line/space features allowed for the cells to migrate in both directions into the gap spacing on each side of the line feature, similar to the migration behavior with a single spacing while still creating an overall unidirectional alignment. Repetitive features allowed for this alignment to continually occur with each line/space repetition, resulting in long-range cellular order perpendicular to the original features (Fig. 1A).

Within the first 2 days after patterning, cell migration was initiated, and the original features were no longer visible. Cellular alignment was established by day 2, and only strengthened as the cells began to form myotubes and mature to day 6. There was clear directional alignment established over the entire patterned substrate, through verification of myosin staining, which was not seen in the negative control condition. This reinforces the fact that we are achieving rapid myotube alignment that is only strengthened as the cells mature into myotubes. We compared common myotube metrics (myotube length, number of nuclei per myotube) across each seeding condition. Similar tube structures were formed for each seeding condition and were comparable to those found using other alignment strategies in the literature (35–39). This indicates that biotinylating the cell surface and conjugating the collagen I substrate with biotin-streptavidin does not alter the intrinsic cell behavior or long-term functionality.

We determined that cell patterning using biotin-streptavidin conjugations creates only a temporary positioning of the cells based on their ability to migrate from their original position. Additionally, the biotin-streptavidin surface patterns were monitored over time via a Cy3 fluorophore tagged to the streptavidin molecules to determine the overall pattern stability on the substrate surface. The biotin-streptavidin surface patterns were confirmed to be temporary as the intensity of the patterns diminished, which can occur due to pH changes in the media or repetitive imaging (40, 41) (Supplementary Fig. 2, 3). This observation further solidifies that the biotin-streptavidin patterning system we have created is only a temporary means for cell placement, which does not pose any immediate or long-term concerns to cell health or function.

By using our biotin-streptavidin patterning system, we achieved rapid cellular alignment and myotube formation over a large 2D surface area. This result was true using primary MPCs from more than one human donor (Fig. 5, Supplementary Fig. 4). The ability to create rapid myotube alignment from multiple donor cell lines signifies that our system is a customizable platform that can be individualized to each patient. Our patterning approach has potential to assist in designing individualized tissue constructs with long-range myotube orientation, with the goal to integrate at the injury site with the existing tissue and reduce the risk of rejection.

## 5. Conclusions

Myotube alignment can be achieved by utilizing the MPC’s innate regenerative response. Biotin and streptavidin molecules provide a rapid means of positioning MPCs in temporary line structures with intentional spacings. These spacings create a small-scale injury model that initiates the MPCs to migrate over the gaps and form myotubes. Biotin-streptavidin conjugations do not impede the intrinsic migratory of MPCs or alter the properties of the myotube structures that they form. Creating repetitions of line/space features using our patterning system allows for long-range ordering of MPCs based on their natural migratory behavior. We have observed this phenomenon to be true using two donor cell lines, showing the potential of our system for widespread use in individualized tissue construct designs.

## 6 Acknowledgements

We would like to thank the University of Kentucky Center for Muscle Biology for their expertise and for providing the human primary myogenic progenitor cells (MPCs) used in these studies.

