## Supplementary Information for "Spontaneous Alignment of Myotubes through Myogenic Progenitor Cell Migration"

**Supplementary Table 1. Significance testing of myotube metrics**

Summary of significance testing using an independent two-group Student's t-test comparing myotube length and the number of nuclei per myotube for myotubes formed from MPCs in each seeding condition (negative control and pattern alignment). Significance was determined as * $p < 0.05$.

| Seeding Condition 1 | Seeding Condition 2 | Variable | P-Value |
|---|---|---|---|
| Negative Control | Pattern Alignment | Myotube Length | 0.47 |
| Negative Control | Pattern Alignment | Nuclei Per Myotube | 0.53 |

SI 1. Cell Migration Using Circular Patterned Structures

Cells were patterned according to the protocol described in Sections 2.3 and 2.4 in 1000 μm diameter circular structures on collagen I to determine any outward cell migration behavior over 6 days in differentiation media. Single circular structures and double circular structures with 800 μm spacing were patterned to determine cell migration towards or away from other cells. Cells were imaged using brightfield microscopy every two days for a total of six days and ImageJ (NIH) Directionality plug-in was used to analyze cell behavior. The slides were fixed on Day 6 with 4% formaldehyde buffer, permeabilized, stained with a DAPI nuclear stain, and imaged for cell counting analysis.

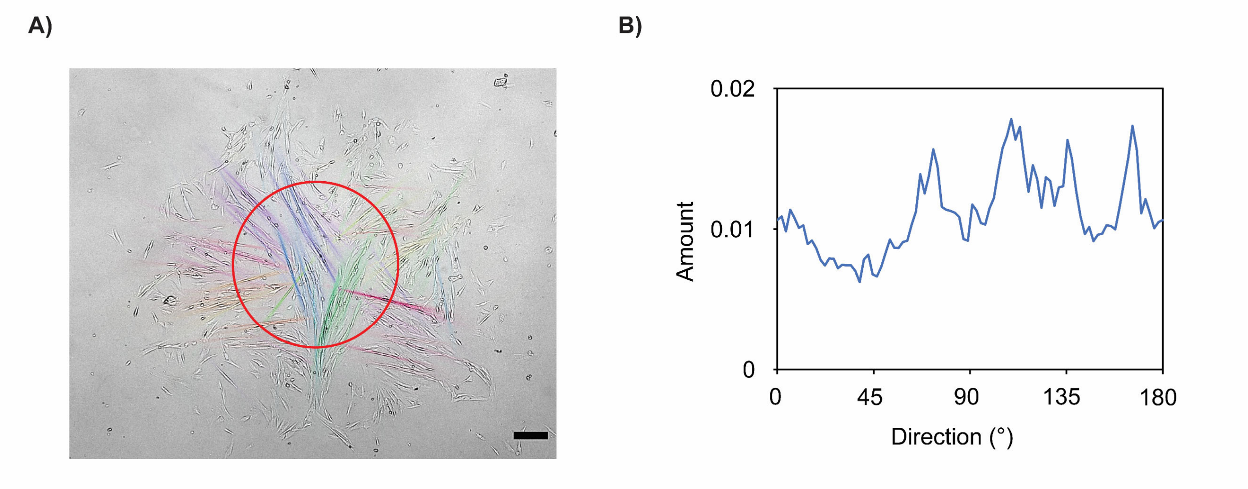


**Supplementary Figure 1. Directionality of single circular pattern feature**
A) Representative brightfield image of MPCs patterned on a 1000 μm diameter circular pattern using biotin-streptavidin conjugations and placed in differentiation media for 2 days. The boundary of the original biotin-streptavidin surface pattern is shown in red. The color overlay on the image represents the various cellular orientations generated by ImageJ.
B) Directionality analysis of MPCs in the image in A.
Scale bars represent 200 μm.

SI 2. Stability of Surface Patterns

The stability of the biotin-streptavidin patterns on the collagen 1 substrate was quantified using a microarray scanner (Affymetrix 428). Patterned slides were scanned with the array scanner and analyzed with ImageJ to determine the pixel intensity within the patterned and unpatterned areas of the slide. 10 data points were collected within four patterned and four unpatterned lines in the middle of the sample. To determine the stability of the patterns over time, a patterned slide was scanned immediately after being patterned, then placed in differentiation media at 37˚ C and 5% $CO_2$. Every 24 h, the slide was removed from the culture conditions, washed with DI $H_2O$ and air dried, and scanned for the pixel intensity. This process was repeated for 10 days with n = 3 independent samples to calculate the sample mean and standard deviation. As a control, a patterned slide was scanned repetitively 10 times to confirm that the array scanner did not cause photobleaching of the patterned structures. Additionally, a patterned slide was repetitively hydrated in differentiation media, then rinsed with DI water, air dried, and scanned to determine if hydration with media changed stability of the patterned structures.

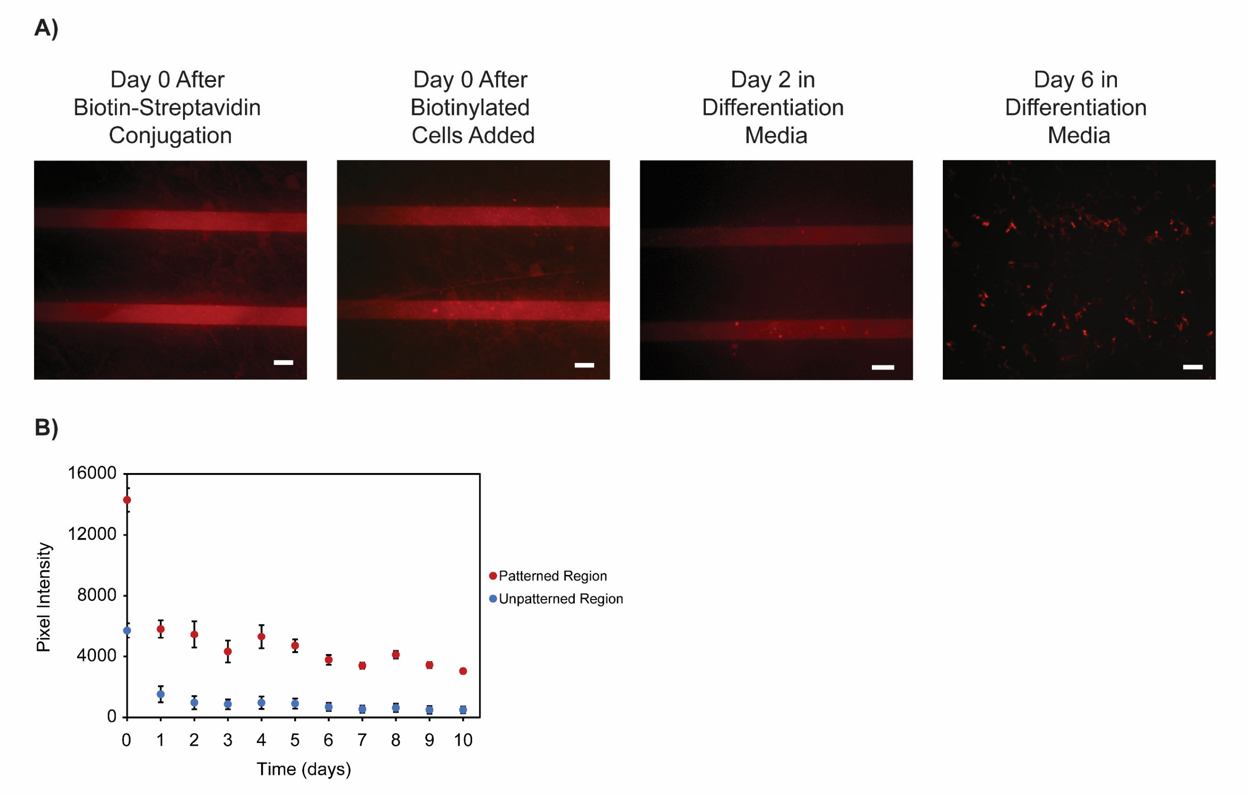


**Supplementary Figure 2. Stability of surface patterns over time.**
A) Representative fluorescent images of biotin-streptavidin surface patterns visualized directly after patterning, the addition of MPCs, and over 6 days in differentiation media.
B) Quantitative analysis of the pixel intensity of patterns over time using the Affymetrix 428 microarray scanner. Scale bars represent 200 µm.

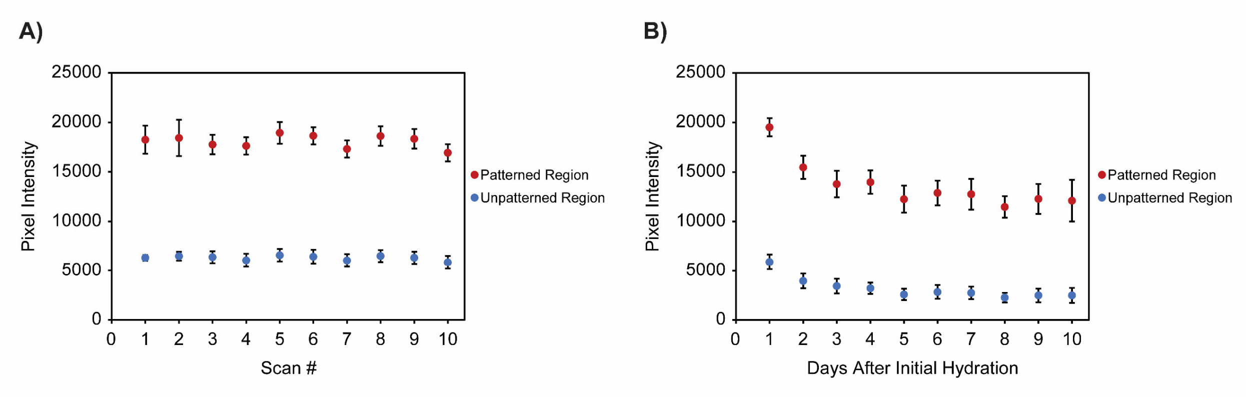


**Supplementary Figure 3. Stability of surface patterns with repetitive hydration and repetitive scanning.**
A) Quantitative analysis of the pixel intensity of patterns with immediate repetitive scanning using the Affymetrix 428 microarray scanner.
B) Quantitative analysis of the pixel intensity of patterns with repetitive hydration in differentiation media using the Affymetrix 428 microarray scanner.

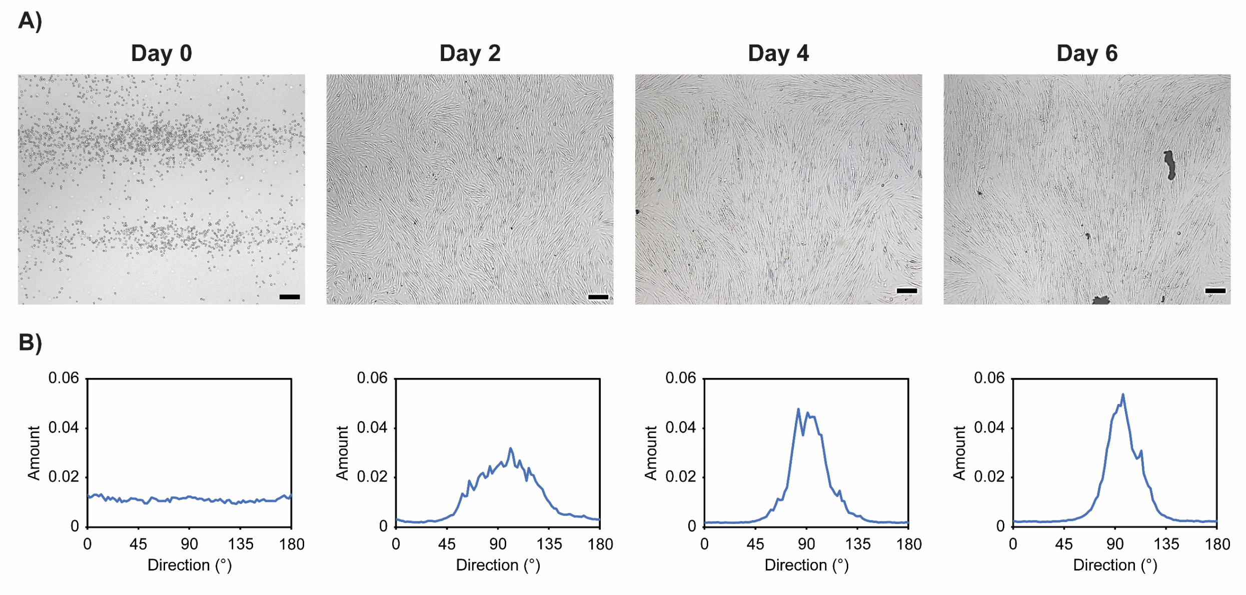


**Supplementary Figure 4. Cell migration analysis of MPCs from second human donor.**
A) Representative images of MPCs placed in temporary patterns of 200 µm line widths with 800 µm spacings using biotin-streptavidin conjugations on Day 0 and observed in differentiation media every 2 days for 6 days. All images were taken using brightfield microscopy at 4x magnification. Scale bars represent 200 µm.
B) Directionality analysis of MPCs at each timepoint mentioned in A.
